# Short-, medium- and long-term metabolic responses of adult ewes submitted to nutritional and β-adrenergic challenges

**DOI:** 10.1101/723965

**Authors:** Eliel González-García, Moutaz Alhamada, Nathalie Debus, Jean-Baptiste Menassol, Anne Tesnière, Jéssica Gonçalves Vero, Bruna Barboza, François Bocquier

## Abstract

**Background:** In order to maintain homeostasis, ruminants submitted to alternating shortage and refeeding situations manifest switches in metabolic pathways induced by undernutrition and body reserves (**BR**) replenishment cycles. The objective of this experiment was to study adaptive regulatory mechanisms present during subsequent feeding transition periods and the inherent lipolytic activity of the adipose tissue in individuals with contrasted BR. Three diets containing different levels of energy were offered to 36 mature, dry, non-pregnant Mérinos d’Arles ewes in an experiment lasting 122 days. Ewes were selected with similar body weight (**BW**), body condition score (**BCS**) and were allocated into three equivalent treatments according to the plane of nutrition: normally fed (**Control**); underfed (**Under**) or overfed (**Over**). The BW, BCS and individual energy metabolism were monitored. At the end of the experiment, lipolytic activity of adipose tissue was studied through a ß-adrenergic challenge to the same ewes, with body conditions according to the offered diet (**Normal, Leans** and **Fat**, respectively).

**Results:** Anabolic or catabolic responses to energy dietary manipulation were accompanied by synchronised metabolic regulation, leading to contrasting metabolic and BR profiles. Average BW and BCS were higher and lower in Over and Under ewes, respectively. The higher and lower BR variations were observed for Under and Over ewes. Higher plasma non-esterified fatty acids (**NEFA**) concentrations were accompanied by lower insulin, leptin and glucose. Differences in leptin were consistent with the dietary energy load (Over > Control > Under). After refeeding, a rebound in BW and BCS was observed for the three groups whereas NEFA was drastically reduced in Under ewes. No differences among treatments were detected in NEFA profiles at the end of the study but lipolytic activity responses to the ß - adrenergic challenge were different and coherent with the adipose tissue mass (Fat > Normal > Lean) and, importantly, was also different between ewes from the same group or BR status, thus evidencing diversity among individual adaptive capacities.

**Conclusions:** The ability of ewes to quickly overcome undernutrition situations by efficiently using their BR was confirmed. There is potential for a simplified ß-adrenergic challenge protocol helping to identify differences in adaptive capacity among individuals.

## Background

Maintaining the consistency of the internal environment (homeostasis) and/or sustaining productive functions (homeorhesis) are essential mechanisms of control in ruminants, allowing them to adapt to physiological and environmental fluctuations [1].

How the animal partitions its nutrients when resources are limited, or imbalanced, is a major way in which it is able to cope with such variations, and thus, determines its robustness. In highly productive selected ruminants there is evidence that their reliance on body reserves (**BR**) is increased and robustness is reduced [2]. The efficiency of BR mobilisation-accretion processes, in order to overcome undernutrition events, is therefore recognised as an essential trait in ruminants. These processes contribute to maintaining the resilience of the flock under fluctuating circumstances, such as in tropical or Mediterranean regions, where seasonal forage availability is highly contrasted.

In previous works characterizing the energy metabolism in a typical round productive year of Romane [3] and Lacaune [4] meat and dairy ewes, respectively, the potential of plasma non-esterified fatty acids (**NEFA**) for being used as predictor of the ruminant nutritional status was confirmed. Furthermore, we know that adipose tissue (**AT**) lipolytic potential can be estimated *in vitro* (by glycerol and NEFA responses from tissue explants into the incubation medium) or *in vivo* by plasma glycerol or NEFA response to injection or infusion of catecholamines or synthetic drugs (β-adrenergic agonists) [5]. Such lipolytic potential could be seen as a sight of the ultimate necessity of the animal to compensate their basic requirements by using their BR. When facing an undernutrition event, a quick BR mobilization (illustrated by plasma NEFA) could be a symptom of the incapacity of the animal to re-adjust its maintenance energy requirements (**MER**) which would lead to regulate (reduce) its feed intake. Under the same conditions (i.e. species, breed, physiological state, age, production system, feeding regimen…) less NEFA in the immediate response would means that the animal is less depending from its BR in the very short term.

For this study we hypothesised that offering restricted diets to adult Mérinos d’Arles ewes would significantly increase their BR mobilization to meeting their MER. After refeeding, the metabolic plasticity of the breed [6,7,8] would lead to recovery within a similar period of time to that of feed restriction. We also hypothesised that those ewes with contrasted body condition scores (**BCS**), resulting from receiving different dietary regimes, would respond differently to an *in vivo* β-adrenergic challenge. That response will correspond to the individual reactivity or adaptive capacity.

Thus, the objective of this study was to evaluate the impact of offering diets of differing nutritional planes on the adaptive capacity of mature ewes at the short-, medium- and long-term. Such adaptation will be characterized by studying trends in the individual BR mobilisation-accretion and the associated metabolic profiles after dietary challenge. A second objective was to evaluate the impact of different BCS on the individual lipolytic potential of the adipose tissue of the ewes facing a β-adrenergic challenge. This would allow us to study the potential of a simplified method for analysing the intraflock variability in individual metabolic plasticity responses when facing nutritional alias.

## Methods

### Location

The experiment was conducted at the Montpellier SupAgro Domaine du Merle experimental farm, located in Salon-de-Provence in the south-east of France (43°38’N, 5°00’E). All animals were cared for in accordance with the guidelines of the Institut National de la Recherche Agronomique (INRA) animal ethics committee.

### Ewes, management, feeding and experimental design

After weaning their litters in mid-January, 36 adult Mérinos d’Arles ewes between 6 and 10 years old and being lambed in October (average lambing on 10 October) were selected for this study from the main research flock. The body weight (**BW**) and BCS was used to select animals with similar body conditions. The initial BW and BCS were 44.4 ± 0.83 kg and 2.0 ± 0.05, respectively.

A schematic representation of the experimental design is presented in Figure 1. The experiment lasted 122 days, and was comprised of two consecutive periods. Firstly, the ewes were allowed to acclimatise to the feeding regimen and the general environment of the sheepfold for 22 days (under confinement). All ewes were managed as a single flock and fed the same Control diet (composition included below) throughout this period. Following acclimatisation was a measurement period of 100 days, beginning from the point at which the experimental feeding regime began (day “zero”). Ewes were randomly assigned to one of three covered pens, each with an area of approximately 30 m^2^ and containing both concrete and straw flooring in the same sheepfold. The 100 day measurement period was stratified into two sub-periods of similar lengths (50 days each), including a dietary challenge period (from day 0 to 49) followed by a refeeding period (from day 50 to 100; Figure 1). At the end of the experiment (last day) a ß-adrenergic challenge protocol was carried out (details included below).

**Figure 1.**
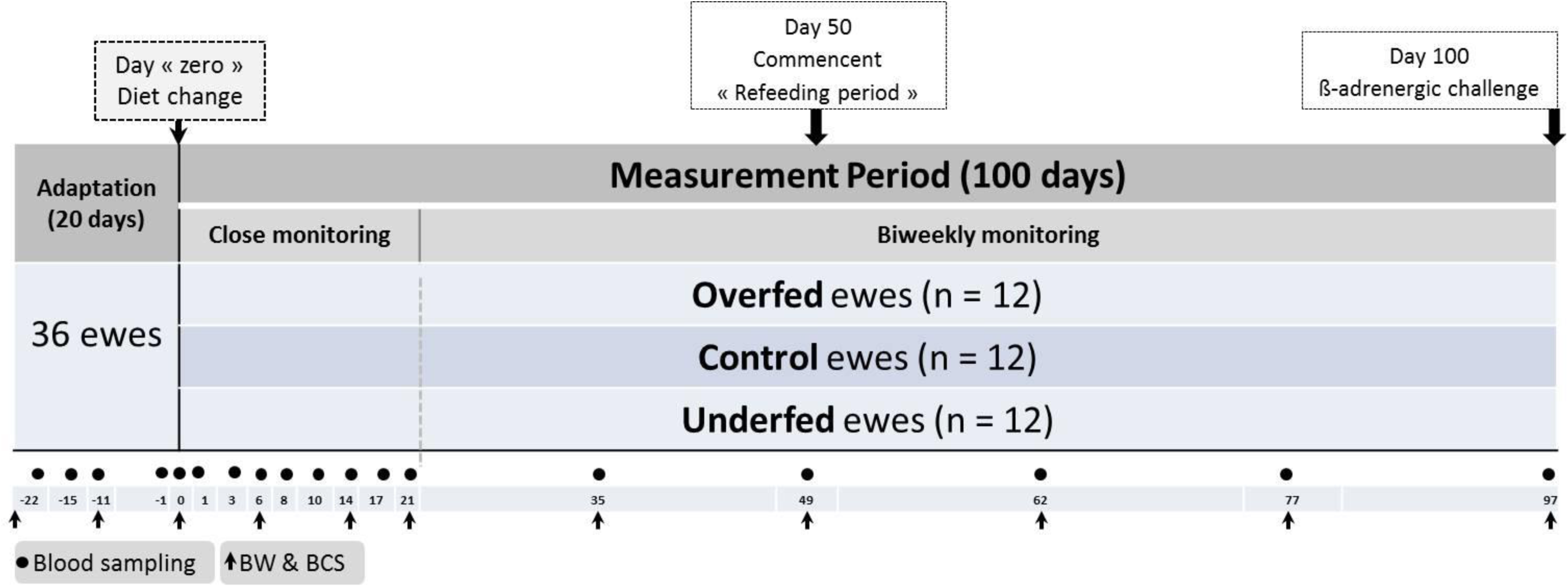
Schematic representation of the experimental design. The distribution of experimental ewes (n = 36) submitted to three contrasted nutritional planes and the body weight (n = 11), blood (n = 18) sampling points and a final ß-adrenergic challenge are illustrated. After 3 weeks of adaptation, energy diet content was changed in the two extreme groups (overfed and underfed). The measurement period (100 days) consisted in two periods i.e. dietary challenge (from 0 to 49 days) and refeeding (from 50 to 100 days) period. Sampling was structured in a close (3 weeks) and a more extended (biweekly) individual monitoring periods.

Ewes were pen-fed and feed was allocated according to treatment to the three groups in each of the covered pens (n = 12 ewes/pen). The treatments included three contrasted diets of different nutrition planes depending on energy supply: i) underfed ewes offered 70% of theoretical MER (i.e. **Under** group), ii) ewes offered 100% of MER (i.e. **Control** group) and iii) ewes offered 160% of MER (i.e. overfed ewes, **Over** group).

At the start of the experiment, ewes were in a maintenance state. Considering the average BW (∼45 kg BW; 17.4 kg BW.^75^) the individual daily intake capacity was 1.3 fill units. According to the INRA tables [9], in order to meet their MER 0.033 fill unit/kg BW.^75^ for maintenance and meat production (**UFV**) and 2.3 g/kg BW.^75^ of protein digestible in the small intestine (**PDI**) was required. Therefore, feeding regimes for each treatment were theoretically planned to achieve three different BCS of 1.75, 2.5 and 3.25 for Under, Control and Over ewes, respectively.

The nutritive values of the ingredients included in the experimental diets are presented in Table 1. As the basal roughage, a wheat straw containing 3.5% of crude protein (CP), 1.34 Mcal/kg dry matter (DM) of metabolisable energy (ME) and 2.4 UFV (INRA, 2010; [9]) was used. Dried and pelleted alfalfa (16% CP) was offered as the main protein source, whereas a dried and pelleted sugar beet pulp was supplied as the main energy source (2.7 Mcal/kg DM of ME and 1.0 fill unit for sheep). A mineral-vitamin premix, containing 90 and 126 g/kg DM of P and Ca, respectively, was supplied at the same dose (∼10 g/ewe/day) for all treatments (Table 2), thus, ensuring the same amount of P and Ca (1 g/ewe/day).

**Table 1.** Nutritive value of ingredients included in the experimental diets

| Ingredient | <sup>1</sup>DM,<br>% | Organic<br>constituents,<br>g/ kg DM |  |  | Energy, Mcal/<br>kg DM |  | Minerals,<br>g/ kg DM |  |  | Net energy, /kg<br>DM |  | Protein value,<br>g/ kg DM |  |  | Fill<br>value |
| --- | --- | --- | --- | --- | --- | --- | --- | --- | --- | --- | --- | --- | --- | --- | --- |
|  |  | <sup>2</sup>OM | <sup>3</sup>CP | <sup>4</sup>CF | <sup>5</sup>GE | <sup>6</sup>ME | Ash | P | Ca | <sup>7</sup>UFL | <sup>8</sup>UFV | <sup>9</sup>PDIA | <sup>10</sup>PDIN | <sup>11</sup>PDIE | <sup>12</sup>SFU |
| Wheat<br>straw | 88 | 920 | 35 | 420 | 4.34 | 1.34 | 80 | 1 | 2 | 0.42 | 0.31 | 11 | 22 | 44 | 2.41 |
| Alfalfa<br>(pelleted) | 91 | 885 | 160 | 310 | 4.40 | 1.84 | 115 | - | - | 0.67 | 0.56 | 50 | 101 | 87 | 0.80 |
| Dried sugar<br>beet pulp | 89 | 912 | 98 | 206 | 4.01 | 2.73 | 88 | 1 | 13 | 1.01 | 0.99 | 40 | 63 | 106 | 1.36 |
| Mineral-vitamin<br>premix | 90 | - | - | - | - | - | - | 90 | 126 | - | - | - | - | - | - |

<sup>1</sup>DM= Dry matter content; <sup>2</sup>OM= Organic matter content; <sup>3</sup>CP= Crude protein content; <sup>4</sup>CF= Crude fibre content; <sup>5</sup>GE= Gross energy; <sup>6</sup>ME= metabolizable energy; <sup>7</sup>UFL= net energy for lactation; <sup>8</sup>UFV= net energy for maintenance and meat production; <sup>9</sup>PDIA= dietary protein undegraded in the rumen which is digestible in the small intestine; <sup>10</sup>PDIN= PDIA + PDIMN (microbial protein that could be synthesized from the rumen degraded dietary N when energy is not limiting); <sup>11</sup>PDIE= PDIA + PDIME (microbial protein that could be synthesized from the energy available in the rumen when degraded N is not limiting); <sup>12</sup>SFU= Fill unit for sheep

**Table 2.** Diet composition and daily nutrient supply according to treatment during the dietary challenge period (day 0 to 49). Values between brackets correspond to the added or reduced quantities applied during the refeeding period (day 49 till 100)

| Treatment | Ingredient<sup>a</sup> | Distributed<br>(as-fed), kg | Nutrient supply per ewe (DM basis) |  |  |  |  |  |  |  |
| --- | --- | --- | --- | --- | --- | --- | --- | --- | --- | --- |
|  |  |  | DM, kg | <sup>1</sup>UFV | <sup>2</sup>PDIN, g | <sup>3</sup>PDIE, g | Fill units | <sup>4</sup>CP, g | P, g | Ca, g |
| Control<br>(100%<br><sup>6</sup>MER) | Wheat straw | 0.91 (=) | 0.80 (=) | 0.25 (=) | 18 (=) | 35 (=) | 1.28 (=) | 28 (=) | 1 (=) | 2 (=) |
|  | Alfalfa | 0.17 (+11) | 0.15 (+0.10) | 0.08 (+0.06) | 15 (+10) | 13 (+9) | 0.12 (+0.8) | 24 (+16) | 1 (+0.5) | 1 (+0.5) |
|  | Dried sugar beet pulp | 0.17 (+11) | 0.15 (+0.10) | 0.15 (+0.10) | 9 (+7) | 16 (+11) | 0.12 (+0.8) | 15 (+10) | 0 (+0.3) | 2 |
| Underfed<br>(70% MER) | Wheat straw | 1.00 (-0.50) | 0.88 (-0.38) | 0.27 (-0.11) | 19 (-8) | 39 (-17) | 1.41 (-0.61) | 31 (-13) | 1 (=) | 2 |
|  | Alfalfa | 0 (+10) | 0 (+0.10) | 0 (+0.06) | 0 (+10) | 0 (+9) | 0 (+0.08) | 0 (+16) | 0 (=) | 0 (=) |
|  | Dried sugar beet pulp | 0 (+11) | 0 (+0.10) | 0 (+0.10) | 0 (+6) | 0 (+11) | 0 (+0.08) | 0 (+10) | 0 (=) | 0 (+1) |
| Overfed<br>(160%<br>MER) | Wheat straw | 0.57 (+11) | 0.50 (=) | 0.16 (=) | 11 (=) | 22 (=) | 0.80 (=) | 18 (=) | 1 (=) | 1 (=) |
|  | Alfalfa | 0.44 (+10) | 0.40 (+0.10) | 0.22 (+0.06) | 40 (+11) | 35 (+9) | 0.32 (+0.08) | 64 (+16) | 1 (+0.5) | 1 (+0.5) |
|  | Dried sugar beet pulp | 0.45 (+11) | 0.40 (+0.10) | 0.40 (+0.10) | 25 (+7) | 42 (+11) | 0.32 (+0.08) | 39 (+10) | 0 (+0.5) | 5 (+1.5) |

<sup>1</sup>UFV= net energy for maintenance and meat production; <sup>2</sup>PDIN= PDIA + PDIMN (microbial protein that could be synthesized from the rumen degraded dietary N when energy is not limiting); <sup>3</sup>PDIE= PDIA + PDIME (microbial protein that could be synthesized from the energy available in the rumen when degraded N is not limiting); <sup>4</sup>CP= Crude protein; <sup>6</sup>MER= maintenance energy requirements. <sup>a</sup>The mineral-vitamin premix was supplied at the same rate for all treatments i.e. 10 g/ewe/d thus providing the same amount of P and Ca (1g/ewe/d and 1 g/ewe/d, respectively)

Table 2 presents the dietary composition for each experimental treatment, including the amounts of each ingredient and the overall daily nutrient supply (per ewe), according to each nutritional plane used in the study. The Control diet was composed of 910 g of wheat straw, 165 g of alfalfa and 170 g dried sugar beet pulp. For the Over group, the quantities of alfalfa and sugar beet pulp were increased (almost tripled) compared to Control, whereas the quantity of wheat straw was reduced by almost half. In contrast, the Under (feed-restricted) ewes were offered only 1 kg of wheat straw daily. These experimental diets corresponded to the dietary challenge period from day 0–49 following acclimatisation (Figure 1). During the second half of the measurement period (refeeding), from day 50 to the end of the experiment (day 100), an equivalent additional daily quantity (DM basis) of 100 g of alfalfa and 100 g of dried sugar beet pulp per ewe were supplied to each of the three experimental groups. In this refeeding period, the quantity of wheat straw supplied to the Under group was reduced to half of that supplied during the 0 to 49 day measurement period (Table 2).

Ewes were group fed once daily at 0800 h, and diets were provided *ad libitum*, which was weekly adjusted at 120% compared to the average intake for previous week. Feed refusals were daily weighted and samples were weekly pooled for further analyses. Ewes in each treatment group had free access to fresh drinking water.

### The ß-adrenergic (isoprotenol) challenge

The contrasting BCS attained among groups at the end of the experimental period allowed to induce a ß-adrenergic challenge to the thirty-six mature, dry, non-pregnant Mérinos d’Arles meat ewes. The objective was to evaluate the lipidyc potential of the AT of the 3 contrasted BCS groups i.e. normal ewes issued from the Control diet (**Normal**, n= 12), underfed or lean ewes (**Lean**, n= 12) and overfed, fatty ewes (**Fat**, n= 12). The previous day of the β-adrenergic challenge the ewes were individually weighed and the BCS estimated.

All ewes (n = 36) were challenged early in the morning (∼0800 h) of the same day (day 100). The challenge consisted on an intravenous injection (4 nmol/kg BW) of isoproterenol (ISO, Isuprel™; Hospira France, 92360 Meudon-La-Fôret, France). Isuprel™ (0.2 mg isoproterenol hydrochloride/mL sterile injection) is a potent nonselective β-adrenergic agonist with very low affinity for α-adrenergic receptors. For individual monitoring of reactions, blood samples (n= 10) were individually drawn from each ewe by jugular venipuncture at -15, -5, 0, 5, 10, 15, 20, 30, 45 and 60 min. relative to the β-adrenergic challenge time.

### Measurements, blood sampling, hormones and metabolite assays

Measurements lasted a total of 122 days, starting with the acclimatisation period (22 days) and continuing throughout the 100 day measurement period (Figure 1). Ewes were individually and manually monitored for BW (n= 11) and BCS [10] at 28 and 11 days prior to the experimental period (-28 and -11, respectively), and at day 0, 6, 14, 21, 35, 49, 62, 77 and 97 after the beginning of the dietary challenge. Similarly, plasma samples for the determination of metabolites and metabolic hormones associated with energy metabolism (n = 18) were taken at 22, 15, 11 and 1 day prior to the experimental period (-22, -15, -11 and -1, respectively), and at day 0, 1, 3, 6, 8, 10, 14, 17, 21, 35, 49, 62, 77 and 97 following the beginning of the dietary challenge.

The close monitoring of ewes (every two or three days) started the day before the dietary changes and lasted until 3 weeks after the beginning of the 100-day measurement period. Following this, approximately two sampling points per month were performed until the end of the experiment (see Figure 1 for details on the experimental design schedule).

For monitoring the energy metabolism progression of each experimental group, individual concentrations of plasma metabolites, including non-esterified fatty acids (**NEFA**), beta-hydroxybutyrate (**β-OHB**) and glucose (**GLU**), and the metabolic hormones insulin (**INS**) and leptin (**LEPT**) were determined according to the protocols described by González-García et al. [3,4]. Blood samples were taken by jugular venipuncture before the first meal (approximately at 0800 h) on each sampling day. Two 9 mL samples were drawn from each ewe (1 tube with 18 IU of lithium heparin per 1 ml blood and 1 tube with 1.2–2 mg of potassium EDTA per 1 ml blood; Vacuette^®^ Specimen Collection System, Greiner Bio-One GmbH, Austria). Samples were immediately placed on ice before centrifugation at 3600 × *g* for 20 minutes at 4°C. The plasma was collected and stored at -20°C in individual identified aliquots (3 µL) for the metabolite and hormone analyses. Plasma NEFA was measured in duplicate using the commercially available Wako NEFA-HR(2) R1 and R2 kit (manufactured by Wako Chemicals GmbH, Neuss, Germany, and distributed by Laboratoires Sobioda SAS, Montbonnot, Saint Martin, France); intra- and inter-assay variations averaged 4.9% and 3.5%, respectively. Plasma GLU concentrations were measured in triplicate using a commercially available glucose GOD-PAP kit (reference LP87809; manufactured and distributed by Biolabo SAS, Maizy, France); intra- and inter-assay variations averaged 2.5% and 2.1%, respectively. Plasma β-OHB were measured in duplicate using the enzymatic method proposed by Williamson and Mellanby [11]; intra- and inter-assay variations averaged 8.8% and 3.3%, respectively. Plasma INS was measured in duplicate using a commercially available RIA kit (Insulin-CT; manufactured by MP Biomedicals–Solon, Ohio, USA and distributed by Cisbio Bioassays, Codolet, France); intra- and inter-assay variations averaged 10.3% and 4%, respectively. Plasma LEPT was quantified using the double-antibody leptin RIA procedures with some modifications as described by González-García et al. [3,4]; average intra- and inter-assay coefficients of variation were 5.4% and 4.8%, respectively.

For the ß-adrenergic challenge, the 9 mL blood samples (1 tube with 1.2-2 mg of potassium EDTA per 1 ml blood) drawn from each ewe at each sampling point of the kinetic (n = 10) were placed immediately on ice before centrifugation at 3,600 × *g* for 20 min at 4°C. Plasma was harvested and stored at -20°C until analyses in individual identified aliquots (3 µL). Concentrations of plasma NEFA were analysed in duplicate, similarly to the procedure above described. Intra- and inter-assay variation for these samples averaged 4.74% and 6.98%, respectively.

### Calculation and statistical analyses

Statistical analyses were performed using Statistical Analysis System package (SAS; v. 9.1.3., 2002-2003 by SAS Institute Inc., Cary, NC, USA) [12]. Data were analysed using the PROC MIXED function with repeated measures. The least square means separation procedure using the PDIFF option in SAS was used and the statistical model was as follows:

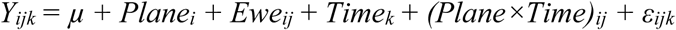

where *Y*_*ijk*_ is the response at time *k*, for ewe *j* that consumed a diet at the nutritional plane *i, µ* is the overall mean, *Plane*_*i*_ is the fixed effect of the specific nutritional plane *i* (_*i*_ = 1–3), *Ewe*_*ij*_ is the random effect of ewe *j* offered the nutritional plane *i, Time*_*k*_ is the fixed effect of time *k, (Plane × Time)*_*ik*_ is the fixed interaction effect of the nutritional plane *i* for time *k* and *ε*_*ijk*_ is the random error at time *k* on ewe *j* offered the nutritional plane *i*.

For the ß-adrenergic challenge database, the NEFA response at each time after challenge was calculated as the change in concentration from basal (-15 min) value as described by Chilliard et al. [13]. The area under the concentration curve (**AUC**) was calculated by doing a definite integral between the two points or limits at time X, using the following formula:

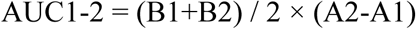

where B is the y axis value (NEFA concentration) and A is the X axis value (time relative to challenge). The AUC was thus calculated for each ewe for the time intervals 0 to 5 min (AUC05), 5 to 10 min (AUC510), 10 to 15 min (AUC1015), 15 to 30 min (AUC1530), 30 to 60 min (AUC3060) and, finally, from 0 to 60 min (AUC060).

By using the data from the concentration-time plot, we calculated the NEFA elimination rate (turnover) constant of each ewe after the ISO challenge (i.e. rate at which NEFA was cleared from the body), assuming first order elimination. For this purpose we calculated K which is the slope of the regression line between time (hours). The measured concentration values of NEFA above the initial point (t = 0; time of injection of ISO) was firstly transformed to their natural logarithm. Extrapolation at zero time gives the theoretical maximal amplitude above initial point (**NEFAamp**). Since we did not have access to the volume diffusion or volume of distribution (V) of NEFA, we used the individual BW of each ewe to determine the clearance rate which was calculated as follows:

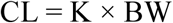

where CL is the NEFA clearance rate value from the body of each ewe, K is the slope of the regression line and BW is the individual BW at the adrenergic challenge moment.

Data of NEFA kinetics in the β-adrenergic challenge were analysed as repeated measures ANOVA using the PROC MIXED function with least squares means separation procedure using the PDIFF option of SAS. The statistical model was as follows:

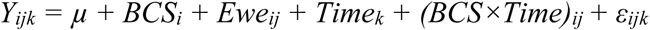

where *Y*_*ijk*_ is the response at time *k* on ewe *j* with a *BCS*_*i*_, *µ* is the overall mean, *BCS*_*i*_ is a fixed effect of the BCS of the ewe at the moment of the challenge *i* (*i* = 1–3), *Ewe*_*ij*_ is a random effect of ewe *j* with a *BCSi, Time*_*k*_ is a fixed effect of time relative to challenge *k, (BCS×Time)*_*ik*_ is a fixed interaction effect of the BCS of the ewe *i* with time relative to challenge *k*, and *ε*_*ijk*_ is random error at time *k* on ewe *j* with a BCS *i*.

The BW, BCS, NEFA responses after challenge with regard to basal NEFA and AUC at different periods were analysed by ANOVA of SAS considering the fixed effect of the BCS group of the observation value. Results were considered significant if *P* < 0.05. Correlation coefficients between basal plasma NEFA and plasma NEFA responses to the ISO challenge and AUC at different ranges of time and from time 0 to 60 min. were determined by using the PROC CORR of SAS.

## Results

The final average individual daily feed balance is shown in Table 3. After calculating the average feed refusal per treatment for each stage of the measurement period (dietary challenge period from 0–49 days and refeeding period from 50–100 days), it was determined that the ewes were 114, 68 and 190% of their MER for the Control, Under and Over groups, respectively. This was different to the 100, 70 and 160% MER theoretically planned, respectively (Table 2). However, the final objective of MER for each of the diets was attained.

**Table 3.**
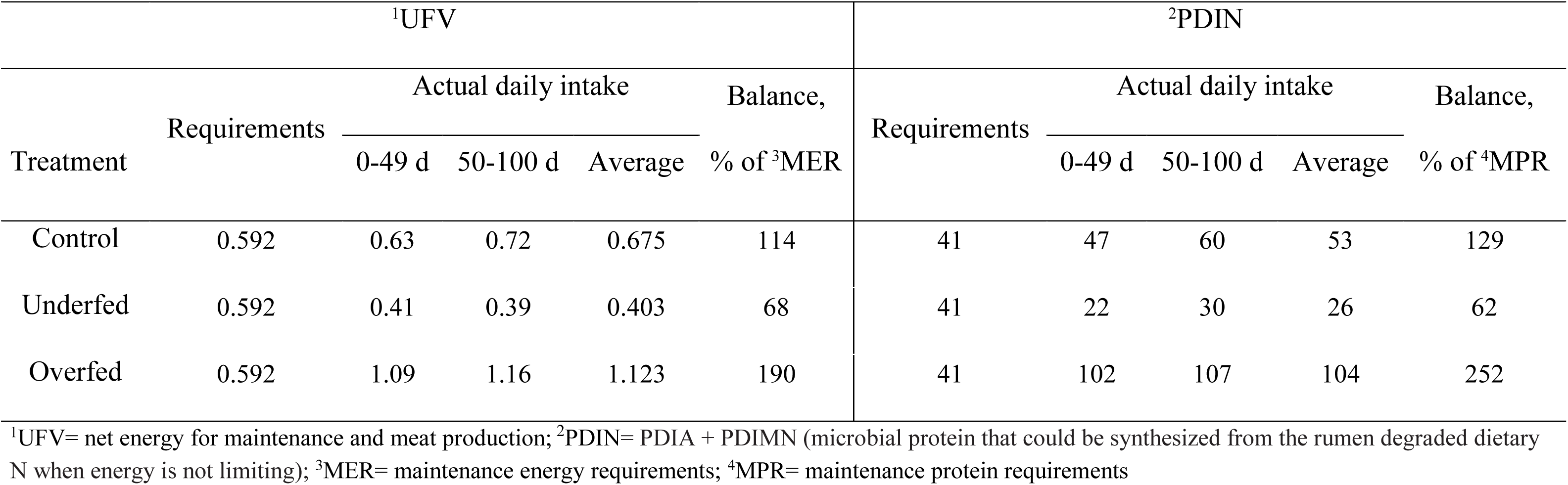
Final individual daily feeding balances, after calculating average feed refusal per treatment

Overall changes in BW, BCS and plasma profiles are presented in Table 4. When all parameters were considered, significant effects were observed for the main sources of variation evaluated i.e. the feeding regimen, time after diet challenge and their first order interactions. A high level of significance was observed for the interaction of nutritional plane with time for all variables measured in this study. As expected, after beginning the feeding regimen (day “zero”) the average BW and BCS were higher and lower, respectively, in the ewes in the Over and Under groups (Table 4 and Figure 2). At the beginning of the refeeding period (day 50) a significant recovery for BW and BCS was observed.

**Table 4.** Average body weight (BW), body condition score (BCS) and plasma profiles of mature, dry, non-pregnant *Mérinos d’Arles* ewes (n = 36) receiving 70% (Underfed; n = 12), 100% (Control; n = 12) or 160% (Overfed; n = 12) of their maintenance energy requirements during the dry-off period

| Item | Nutritional plane or treatment |  |  | Effects, *P* value |  |  |
| --- | --- | --- | --- | --- | --- | --- |
|  | Control | Underfed | Overfed | Regimen | Time | Regimen × Time |
| BW, kg | 44.27 ($\pm 0.285$) | 41.59 ($\pm 0.285$) | 47.48 ($\pm 0.446$) | <.0001 | <.0001 | <.0001 |
| BCS, 1-5 | 1.92 ($\pm 0.017$) | 1.78 ($\pm 0.017$) | 2.00 ($\pm 0.027$) | <.0001 | <.0001 | <.0001 |
| NEFA, mmol/L | 0.12 ($\pm 0.011$) | 0.26 ($\pm 0.011$) | 0.09 ($\pm 0.017$) | <.0001 | 0.0002 | <.0001 |
| β-OHB, mg/L | 23.36 ($\pm 0.684$) | 20.16 ($\pm 0.684$) | 23.58 ($\pm 1.068$) | 0.003 | <.0001 | <.0001 |
| Glucose, g/L | 0.55 ($\pm 0.005$) | 0.55 ($\pm 0.005$) | 0.59 ($\pm 0.008$) | 0.025 | <.0001 | 0.0061 |
| Insulin, μIU/mL | 12.98 ($\pm 0.520$) | 11.58 ($\pm 0.520$) | 14.96 ($\pm 0.813$) | 0.004 | 0.0033 | 0.0001 |
| Leptin, ng/mL | 4.56 ($\pm 0.058$) | 4.52 ($\pm 0.058$) | 5.09 ($\pm 0.091$) | 0.003 | <.0001 | <.0001 |

**Figure 2.**
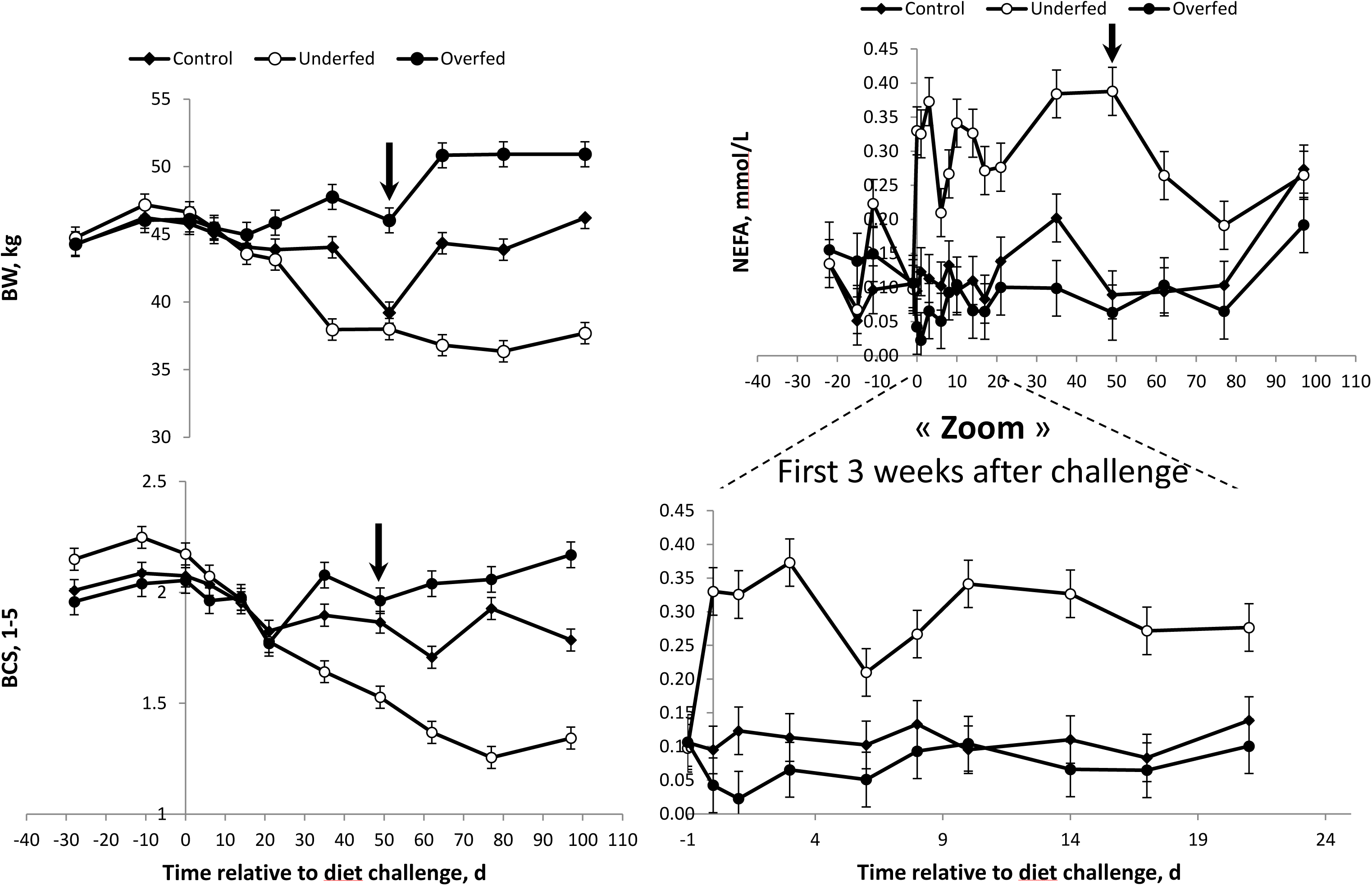
Body weight (BW), body condition score (BCS) and non-esterified fatty acids (**NEFA)** blood plasma profiles of mature, dry, non-pregnant *Mérinos d’Arles* ewes (n = 36) offered 60% (Underfed; n = 12), 100% (Control; n = 12) or 170% (Overfed; n = 12) of maintenance energy requirements. Diet challenge started at day 0, after an overall 3 week adaptation. Arrow represents commencement of refeeding period. Error bars represent SEM.

The differences and trends observed for BW and BCS were consistent with those obtained for NEFA profiles (Figure 2). The higher and lower average BR mobilisation, as illustrated by plasma NEFA concentration, were observed in the Under and Over group of ewes (0.26 ± 0.011 vs. 0.09 ± 0.017 mmol/L, respectively). However, the differences between the Over and Control groups were only evident until 1 week following the change of the feeding regimen (Figure 2). Once started the refeeding period, plasma NEFA was drastically reduced in the Under ewes. At the end of the study, no significant differences were detected between the groups of ewes, regardless of the feeding regimen (Figure 2).

Differences in GLU were only observed when the Over ewes were compared to the other two experimental treatments. However, there were no differences observed when the Under and Control ewes were compared (Table 4 and Figure 3). Conversely, plasma INS concentrations were more consistent with the differences in BR mobilisation rates (i.e. NEFA profiles) shown above. At higher plasma NEFA concentrations a lower INS concentration was obtained. Therefore, plasma INS was higher in the Over ewes, followed by the Control ewes, and was the lowest for ewes in the Under group (Table 4 and Figures 2 and 3). A concomitant, parallel effect on INS and GLU was observed following refeeding. The peak in GLU concentration observed in the Over ewes was followed by the observation of a similar peak for the plasma INS profile at the same time point and in the same group of ewes. In general, either the plasma GLU or INS profiles were higher in the Over ewes throughout the experiment, but no differences were observed when the ewes offered the Control or Under diets were compared (Figure 3).

**Figure 3.**
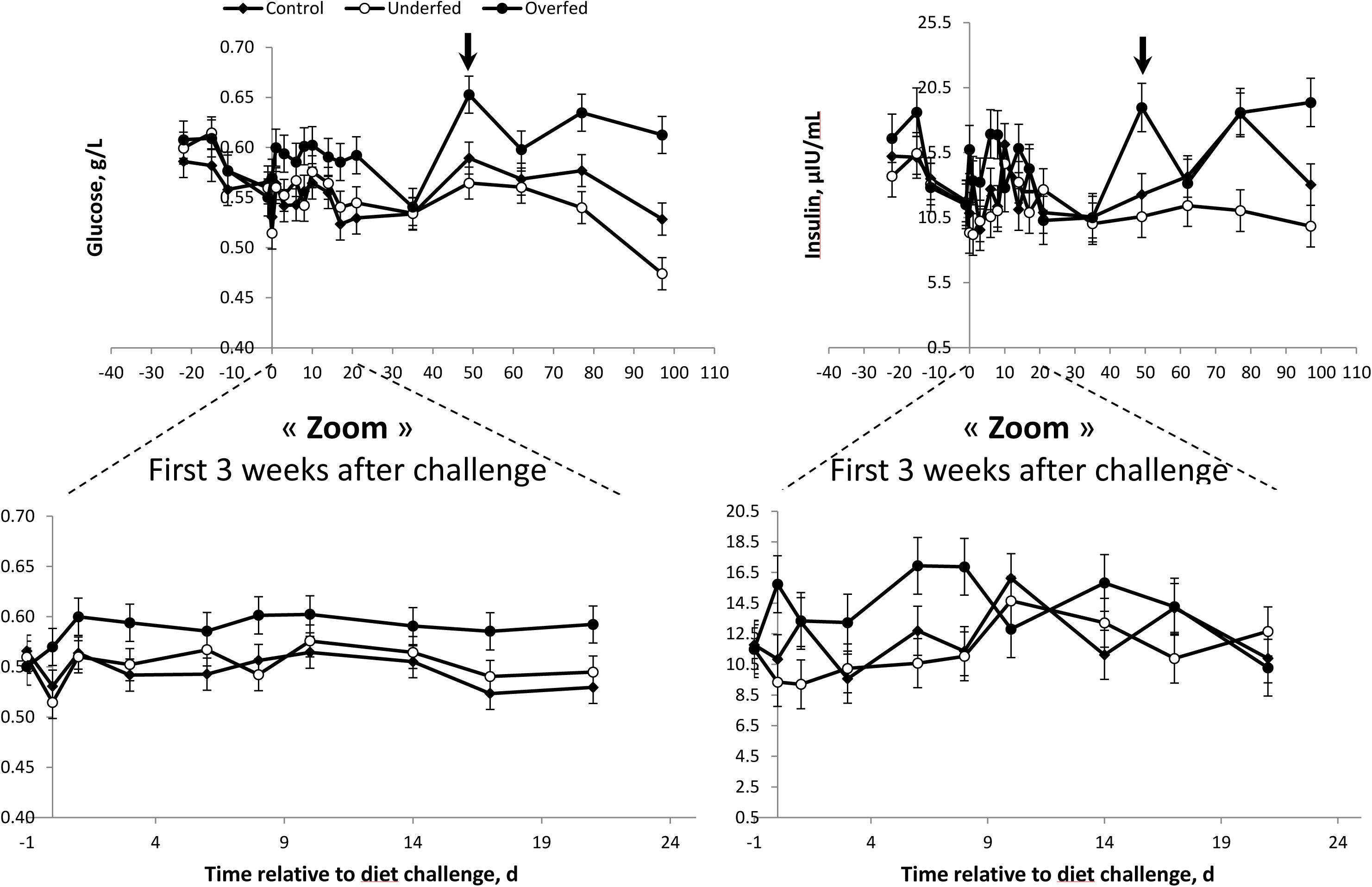
Energy metabolism (**glucose** and **insulin** blood plasma profiles) of mature, dry, non-pregnant *Mérinos d’Arles* ewes (n = 36) offered different nutritional planes during the dry-off period i.e. 60% (Underfed; n = 12), 100% (Control; n = 12) or 170% (Overfed; n = 12) of maintenance energy requirements. Diet challenge started at day 0, after an overall 3 week adaptation period. Arrow represents commencement of refeeding period. Error bars represent SEM.

No differences in ß-OHB concentrations were found when the Control and Over ewes were compared (Table 4). A lower (*P* < 0.003) ß-OHB profile was observed in the Under ewes compared to the average of Control and Over ewes combined (20.16 ± 0.684 vs. 23.47 ± 0.876 mg/L, respectively). The lower ß-OHB profile observed in the Under ewes was consistent throughout the experiment (Figure 4).

**Figure 4.**
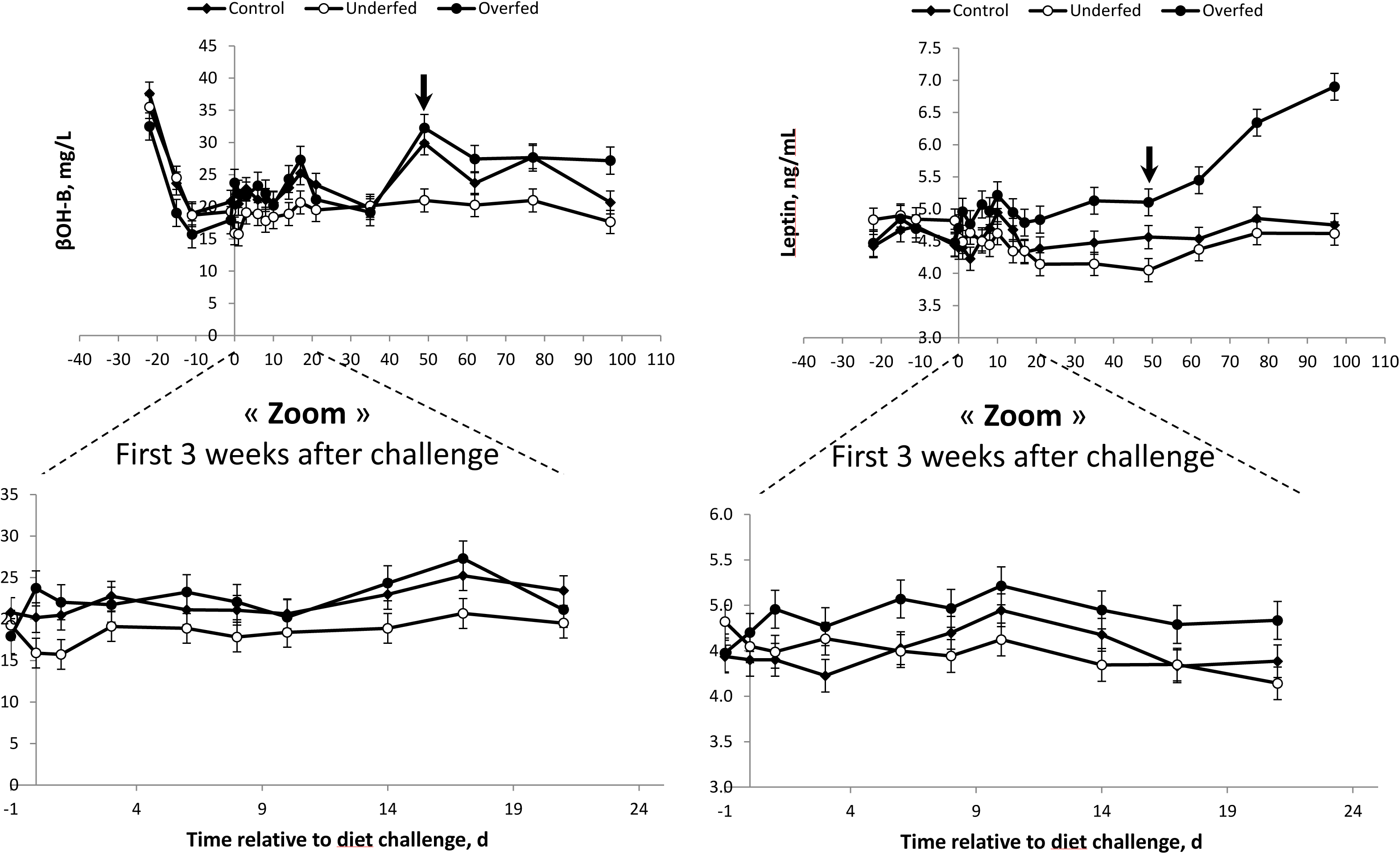
Energy metabolism (β-hydroxybutyrate -**β-OHB**- and **leptin**) blood plasma profile) of mature, dry, non-pregnant *Mérinos d’Arles* ewes (n = 36) offered different nutritional planes during the dry-off period i.e. 60% (Underfed; n = 12), 100% (Control; n = 12) or 170% (Overfed; n = 12) of maintenance energy requirements. Diet challenge started at day 0, after an overall 3 week adaptation period. Arrow represents commencement of refeeding period. Error bars represent SEM.

Differences in plasma LEPT were also consistent with the feeding regimen (Over > Control > Under; Table 4). As expected, a higher LEPT profile was observed in the Over group throughout the experimental period, and these differences were significantly increased following refeeding (Table 4 and Figure 4).

At the day of the ß-adrenergic challenge (end of the experiment), significant differences (*P* < 0.0001) were verified when comparing average BW and BCS of the 3 experimental groups (Table 5; Figure 5). As a result of the previous 100 days feeding manipulation period, ewes belonging to the underfed (Lean) group (BW = 37.7 kg; BCS = 1.34) were more than 10 kg lighter than Normal (BCS = 1.79) and overfed (Fat) (BCS = 2.17) ewes (46.2 and 50.9 kg BW, respectively).

**Table 5.** Average body weight (BW), body condition score (BCS), basal plasma NEFA (at -15 min.) and plasma NEFA responses to a β-adrenergic challenge with isoproterenol injection in mature, dry, non-pregnant *Mérinos d’Arles* ewes (n = 36) with different body condition scores

| Item | BCS group |  |  |  |  |  | Effect, P < |
| --- | --- | --- | --- | --- | --- | --- | --- |
|  | Normal |  | Lean |  | Fat |  |  |
|  | LSmeans | SEM | LSmeans | SEM | LSmeans | SEM |  |
| BW, kg | 46.22 | 0.786 | 37.69 | 0.786 | 50.92 | 0.923 | <.0001 |
| BCS, 1-5 | 1.79 | 0.049 | 1.34 | 0.049 | 2.17 | 0.058 | <.0001 |
| Basal NEFA, mmol/L | 0.06 | 0.02 | 0.11 | 0.02 | 0.05 | 0.02 | 0.0002 |
| Response at 5 min., mmol/L | 0.31 | 0.04 | 0.33 | 0.04 | 0.37 | 0.04 | 0.0003 |
| Response at 10 min., mmol/L | 0.23 | 0.04 | 0.42 | 0.04 | 0.22 | 0.04 | <.0001 |
| Response at 20 min., mmol/L | 0.08 | 0.03 | 0.30 | 0.03 | 0.05 | 0.03 | <.0001 |
| Maximal response, mmol/L | 0.38 | 0.04 | 0.56 | 0.06 | 0.42 | 0.03 | <.0001 |
| Time<sup>¶</sup>, min. | 5.83 | 0.54 | 12.00 | 1.93 | 5.00 | 0.00 | <.0001 |
| AUC<sup>ø</sup>, mmol.min/L |  |  |  |  |  |  |  |
| from 0 to 5 min. | 1.1 | 0.14 | 1.4 | 0.23 | 1.2 | 0.09 | <.0001 |
| from 5 to 10 min. | 1.6 | 0.22 | 2.4 | 0.33 | 1.7 | 0.14 | <.0001 |
| from 10 to 15 min. | 1.2 | 0.20 | 2.6 | 0.31 | 1.1 | 0.12 | <.0001 |
| from 15 to 30 min. | 1.9 | 0.38 | 5.6 | 0.62 | 1.4 | 0.23 | <.0001 |
| from 30 to 60 min. | 2.8 | 0.58 | 5.5 | 0.65 | 2.0 | 0.23 | <.0001 |
| from 0 to 60 min. | 5.4 | 0.87 | 13.3 | 1.53 | 5.5 | 0.59 | <.0001 |

<sup>¶</sup>Time between isoproterenol challenge and maximal response; <sup>ø</sup>Area under the concentration curve and above baseline

**Figure 5.**
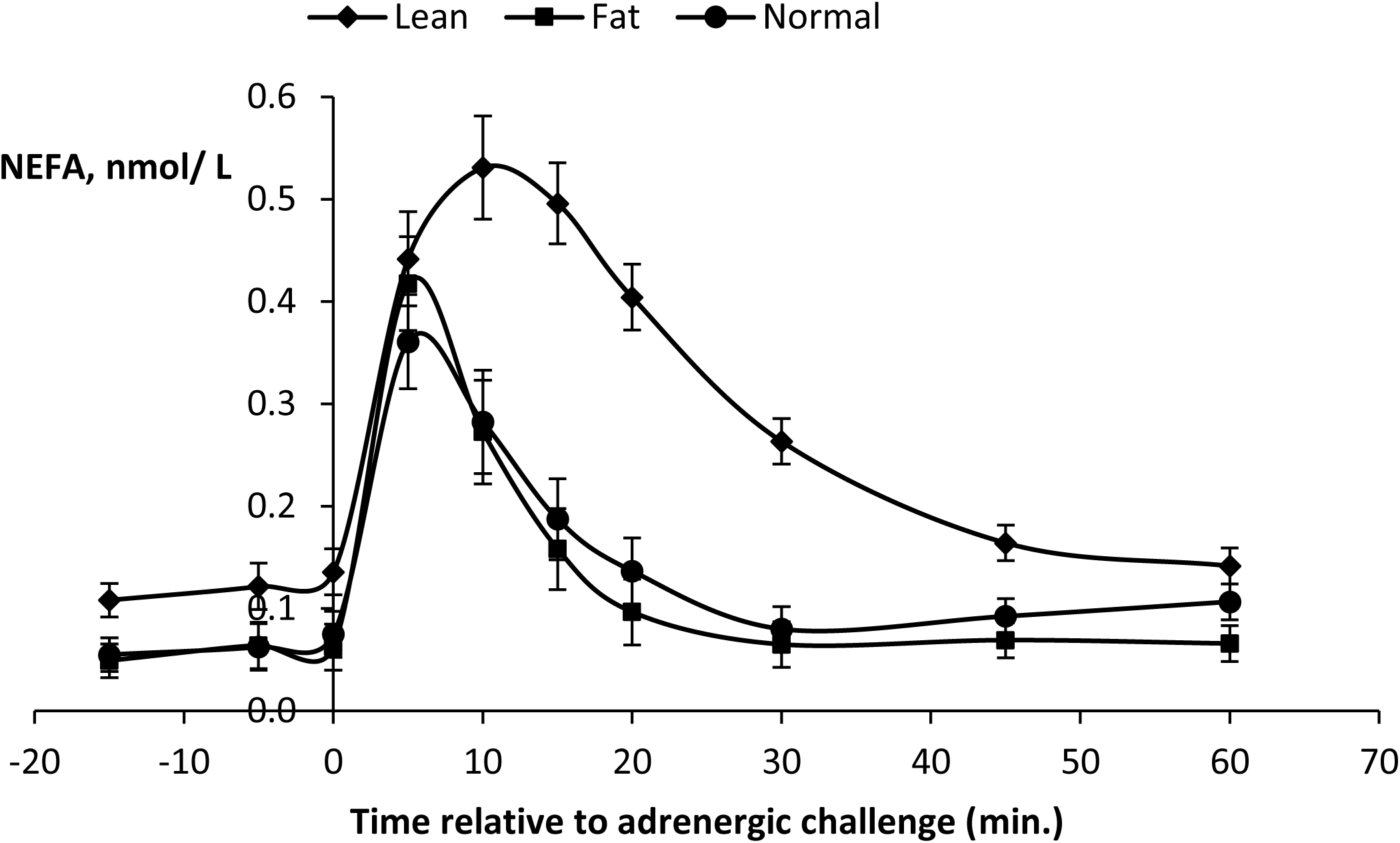
Effect of an induced β-adrenergic challenge injection with isoproterenol (ISO, 4 nmol/kg BW) on plasma non-esterified fatty acids (NEFA) kinetics of mature, dry, non-pregnant *Mérinos d’Arles* ewes (n = 36) with contrasted body condition scores i.e. high (FAT, +BCS; n = 12), average (NORMAL, BCS; n = 12) or low (LEAN, -BCS; n = 12). Error bars represent SEM.

Basal plasma NEFA (–15 min.) before the β-adrenergic challenge was higher (*P* < 0.0002) in Lean ewes compared to Normal and Fat groups (Table 5). In contrast, plasma NEFA response at 5 min. after ISO challenge was higher (*P* < 0.0003) in Fat ewes. After 10 min. plasma NEFA response was consistently higher in Lean ewes. Plasma NEFA maximal response (0.56 mmol/L) was higher (*P* < 0.0001) in Lean ewes and this occurred at 12 min. after the challenge (Table 5; Figure 5).

The AUC were all higher (*P* < 0.0001) in Lean ewes. Thus, overall results showed that underfeeding increased basal plasma NEFA, plasma NEFA response at 10 and 20 min., plasma NEFA maximal response after ISO challenge and all NEFA response areas (*P* < 0.0001). The NEFA maximal response occurred later in underfed or Lean ewes when compared to Control and Over ewes.

The plasma NEFA kinetics for the three experimental groups appears in Figure 5. The NEFA concentrations increased for 10 min. in all groups and were always higher (*P* < 0.0001) in Lean ewes during the 60 min. post-challenge with a peak of plasma NEFA concentration attaining 0.53 mmol/L. However, for all groups NEFA concentration decreased in a similar way and, after 60 min., it returned to values close to baseline.

All correlations (*r* = 0.54 to 0.79) between basal NEFA and different parameters of NEFA response to ISO challenge were significant (*P* < 0.0001; Table 6) with the exception of the variable time between ISO challenge and maximal response (*r* = 0.25). The highest correlations between basal plasma NEFA and responses at different points after challenge were at 10, or 15 min (*r* = 0.73 and 0.69, respectively). The highest correlation between plasma NEFA and AUC were at the area from 0 to 15 min (*r* = 0.79; 0.72 and 0.73 for 0 to 5, 5 to 10 and 10 to 15 min, respectively).

**Table 6.** Correlation coefficients* between basal plasma NEFA and plasma NEFA responses to isoproterenol challenge in fat, lean or normal body condition score meat ewes (n = 36)

|  | Response<br>at<br>5 min | Response<br>at<br>10 min | Response<br>at<br>15 min | Response<br>at<br>20 min | Maximal<br>response | Time<sup>‡</sup> | AUC<sup>∅</sup><br>0-60 | AUC<br>0-5 | AUC<br>5-10 | AUC<br>10-15 | AUC<br>15-30 | AUC<br>30-60 |
| --- | --- | --- | --- | --- | --- | --- | --- | --- | --- | --- | --- | --- |
| Basal NEFA | 0.61 | 0.73 | 0.69 | 0.66 | 0.54 | 0.25 | 0.69 | 0.79 | 0.72 | 0.73 | 0.65 | 0.67 |
| Response at<br>5 min |  | 0.77 | 0.63 | 0.51 | 0.84 | -0.08 | 0.67 | 0.96 | 0.92 | 0.72 | 0.52 | 0.44 |
| Response at<br>10 min |  |  | 0.91 | 0.85 | 0.83 | 0.30 | 0.91 | 0.83 | 0.96 | 0.98 | 0.84 | 0.72 |
| Response at<br>15 min |  |  |  | 0.97 | 0.77 | 0.43 | 0.99 | 0.70 | 0.84 | 0.98 | 0.98 | 0.82 |
| Response at<br>20 min |  |  |  |  | 0.72 | 0.47 | 0.97 | 0.61 | 0.75 | 0.93 | 1.00 | 0.89 |
| Maximal<br>response |  |  |  |  |  | -0.07 | 0.80 | 0.84 | 0.89 | 0.82 | 0.73 | 0.69 |
| Time |  |  |  |  |  |  | 0.40 | 0.01 | 0.15 | 0.37 | 0.47 | 0.44 |
| AUC<br>0-60 |  |  |  |  |  |  |  | 0.74 | 0.86 | 0.97 | 0.97 | 0.83 |
| AUC<br>0-5 |  |  |  |  |  |  |  |  | 0.94 | 0.79 | 0.61 | 0.59 |
| AUC<br>5-10 |  |  |  |  |  |  |  |  |  | 0.92 | 0.75 | 0.64 |
| AUC<br>10-15 |  |  |  |  |  |  |  |  |  |  | 0.93 | 0.79 |
| AUC<br>15-30 |  |  |  |  |  |  |  |  |  |  |  | 0.89 |

\*All significant at $P < 0.0001$ except for the variable Time
<sup>‡</sup>Time between isoproterenol challenge and maximal response; <sup>∅</sup>Area under the concentration curve and above baseline

From 20 min. after challenge, correlations were progressively lower in the sense of the declining tendency of the curve. Correlations with AUC from time 0 to 60 min. were very high for response at 10, 15, 20 min. (*r* = 0.91 to 0.99) and maximal response (*r* = 0.80). Correlations between AUC from time 0 to 60 min. and AUC from time 10 to 15 min., or AUC from time 15 to 30 min. were very high (*r* = 0.97); correlations between AUC from time 0 to 60 min. and AUC from time 30 to 60 min. (declining part of the curve) were also high (*r* = 0.83; Table 6). Correlations between the maximal response and the response at 5, or 10 min., and with the AUC from time 5 to 10 min. were high (ranging from 0.83 to 0.89; Table 6).

Within BCS groups’ correlations between AUC from time 0 to 60 min. and responses at 10, 15 and 20 min were higher than those with basal plasma NEFA, or NEFA response at 5, 30, 45 and 60 min. (Figure 6). Differences in the releasing NEFA turnover were observed between BCS groups and among individuals in the same group and with similar BW and BCS status (Figure 7).

**Figure 6.**
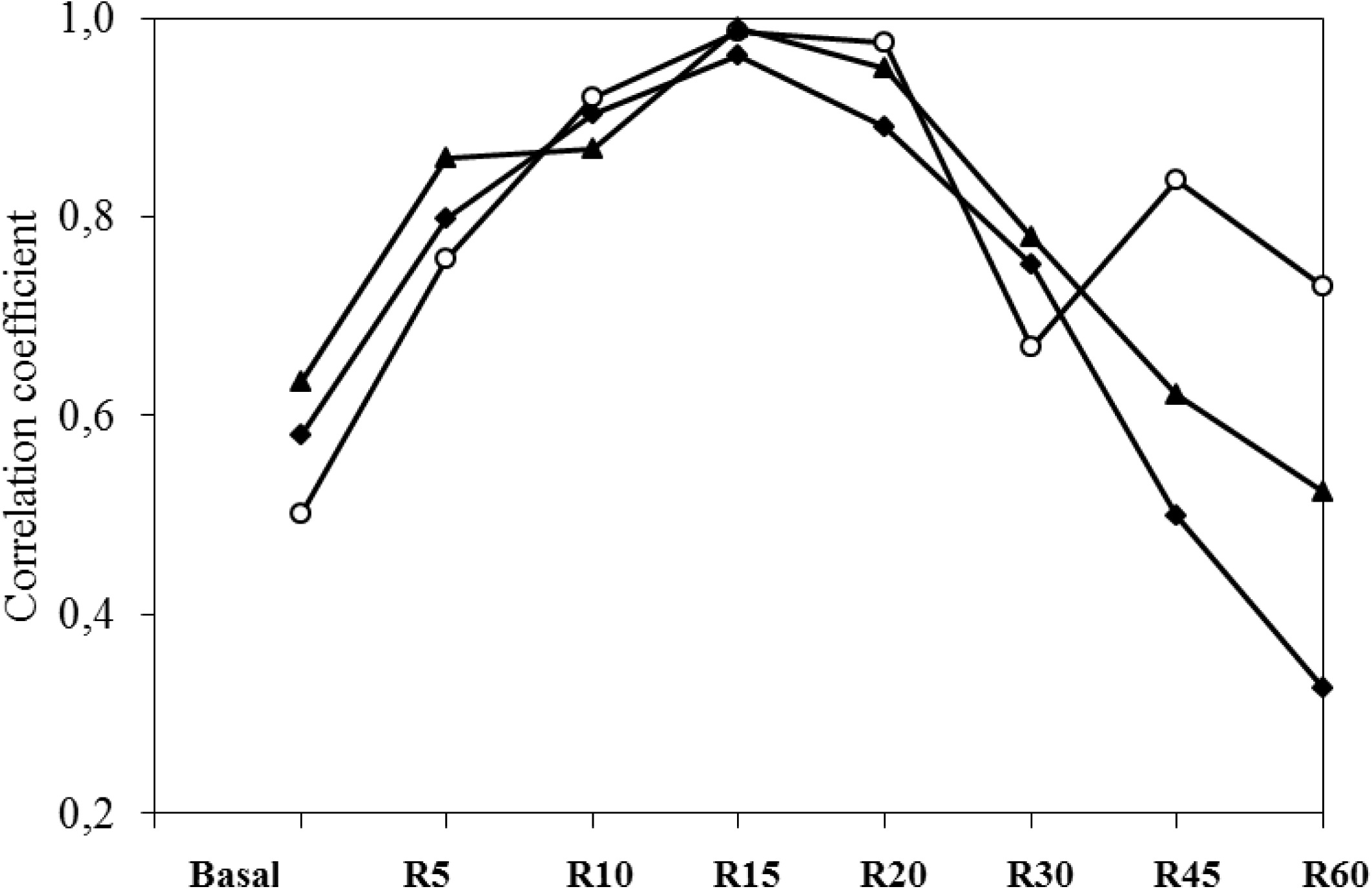
Correlations between plasma area under the concentration curves (AUC) from time 0 to 60 min. and basal plasma NEFA, or plasma NEFA responses at different times after isoproterenol challenge in normal (-°-), lean (-▴) or fat (-•-) *Mérinos d’Arles* meat ewes.

**Figure 7.**
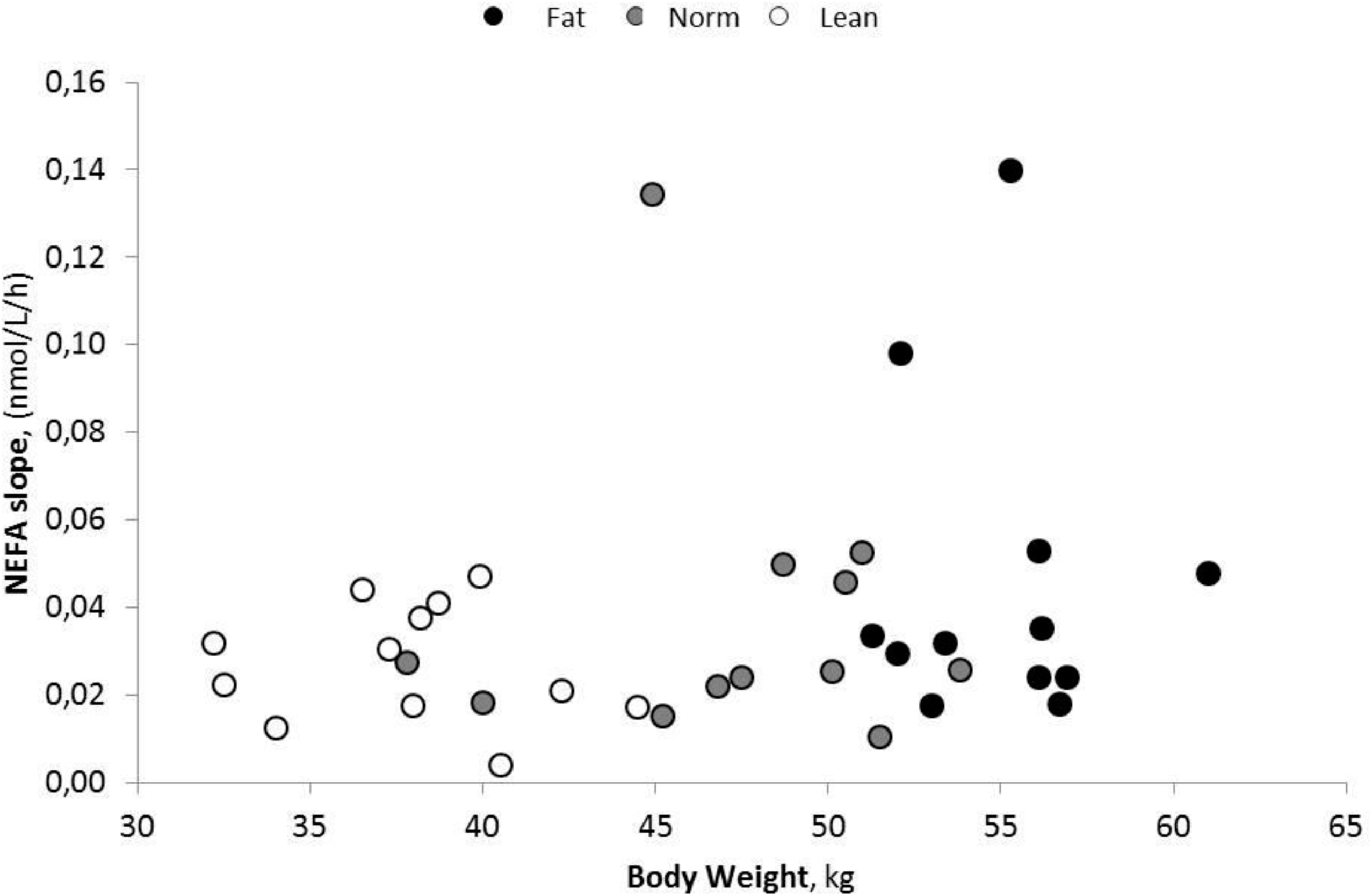
Individual distribution of non-esterified fatty acids (NEFA) release turnover in *Mérinos d’Arles* meat ewes with contrasted body weights (Fat, Normal or Lean) resulting from different dietary challenges (overfed, normally fed or underfed, respectively). Notice that, at similar body weights, ewes belonging to the same group showed contrasted adaptive capacity as illustrated by their short-term responses after a ß-adrenergic challenge.

## Discussion

In the next future, sustainability of farming systems will rely on their ability to cope with a reduction of inputs usage (i.e. concentrate, irrigation, fertilizers…). In this context, a better understanding of the relationship between nutrients supply, nutritional status, their interactions with BR dynamics and the progression of the metabolic profile is essential for the development of a more comprehensive management approach of nutrition based on adaptive capacity of ruminants [14,15,16]. The objectives of this study were to evaluate and describe how dietary energy restriction and/or repletion influence changes in BW, BCS and metabolic status responses in Mérinos d’Arles ewes, considered to be a robust (rustic) and hardy sheep breed.

We validated the previous estimation of energy requirements (INRA, 2007 [9]) of the dry ewe by a stabilized BCS and BW over the whole experimental period with the Control diet. Hence adaptive capacity of ewes fed Under and Over diets will be discussed by direct comparison with the ewes Control responses. We confirmed that offering restricted diets to ewes would induce significant increases in BR mobilisation in order to meet their energy requirements. We also verified that, after partial refeeding, the metabolic plasticity of ewes of this breed allowed the BW and BCS to start recover within a period of similar duration to that of the feed restriction, whereas a high restriction of nutrient allowance would temporarily and negatively affect the voluntary feed intake. The highest feed refusal rate was consistently observed in the Under group, mainly after the first 3–4 weeks of feeding of the restricted diet (data not shown). This is likely to be a consequence of a depressed ruminal environment together with low roughage quality, which is known to negatively affect digestion, metabolism and appetite in undernourished ruminants.

Differences in BW and BCS progression throughout the experiment were expected, considering the dietary energy manipulation. The Under, Control and Over ewes reduced, maintained and improved their BW and BCS, respectively. Ewes in the Under and Control groups responded to the dietary manipulation by attempting to maintain their BW and BCS, as observed with the reduced concentrations of GLU, INS and LEPT, compared to ewes that consumed the Over diets (Table 4 and Figures 2, 3 and 4).

Throughout the experiment, the Under ewes (who consumed almost half of their MER) presented lower BW and BCS, increased plasma NEFA concentration and lower INS and LEPT concentrations when compared to adequately fed animals. The BR mobilisation status was well illustrated by the consistently higher plasma NEFA concentrations observed in Under ewes. This was expected as this group was exposed to a strong dietary energy restriction, based on *ad libitum* wheat straw (containing 3.5 % CP and 1.3 Mcal/kg DM of ME) during the first 50 days of the measurement period. After beginning the partial refeeding period at day 50, the same Under ewes responded to the energy repletion with an immediate short-term decrease in their NEFA concentrations, with more delayed recovery of BW and BCS observed in this group (Figure 2), which is logical since it was only a partial refeeding.

The endocrine system, characterised by plasma INS and LEPT profiles in this study, regulates metabolism by finely tuned peripheral information, which are ultimately aimed to maintain homeostasis. These adaptive processes involve the interplay between several hormones, which also include growth hormone (GH) and insulin-like growth factors (IGFs) [17, 18, 19, 20], which were not analysed in this study.

The typical characteristics of undernourished ruminants were observed in the Under ewes. The lack of glucose arriving to the rumen, in addition to a reduction in volatile fatty acids (VFA) production, induces gluconeogenesis accompanied by intense lipolysis, proteolysis and ketogenesis [1,17]. The reduction of gut metabolic activity is known to account for the decrease of energy requirement in underfed ruminants. Thus, the reduced oxidative or basal metabolism is characterised by a decrease in plasma GLU, INS, LEPT and prolactin concentrations, and an increase in other hormones such as GH, adrenalin, cortisol and glucagon (not measured in this study). This generally leads to shifts in metabolic pathways which aim to spare GLU (with the accompanied increase in NEFA) and proteins (with increased proteolysis and ketogenesis).

Interestingly, the plasma level of β-OHB was lowest in the Under group. This may be due to limits in the supply of β-OHB precursors by this diet. Thus, even if higher BR mobilisation was present in Under ewes, the unexpected lower β-OHB plasma concentration is probably the consequence of the ingredients used in the experimental diets. The Under diet, which only offered wheat straw, did not contain the required precursors. However, with similar underfeeding situation [21] fat-tailed Barbarine ewes were able to produce and survive thanks to their significant ability to mobilise their BR. Plasma NEFA and β-OHB concentrations were initially almost doubled. The medium-term response of these ewes was very similar to what we observed: a steady decline of these metabolites, which was attributed to their ability to adjust their lipid metabolism in order to reduce the toxic effects of high concentrations of NEFA and β-OHB, and therefore, prolong survival. After partial refeeding Barbarine ewes were able to fully recover their initial BW, lipid and protein masses [21].

Submitted to an opposite nutritional situation, the enhancement of the anabolic pathway response was clear in the Over ewes, with increased GLU, INS and LEPT concentrations and decreased plasma NEFA (Figures 2, 3 and 4). However, differences toward dominant anabolic responses were not always evident when comparing the Control and Over ewes, despite the clear differences in energy supply, which should have been sufficient to create greater differences than those observed. However, the BW and BCS progression were clearly different between the groups from 20 days after the introduction of the diet changes (Figure 2). Thus, the differences in responses between these two groups were not well described in this study from the analysis of the chosen metabolites and metabolic hormones. This led us to hypothesise that measuring other parameters, including GH, IGF-1 and polyunsaturated nitrogen (PUN), may provide an improved characterisation of the anabolic and catabolic responses. It is highly probable as reported by Delavaud et al. [22], that increasing amounts of stored body fat in the Over group, increased their MER thus reducing the energy balance gaps between Normal and Over ewes. Hence metabolic and endocrine profiles are blunted by this phenomenon. Such effects of body fatness on energy requirement were reported by Caldeira and Portugal [20] in underfed fat ewes. Ewes with low BCS had lower plasma GLU, triiodothyronine and thyroxine concentrations and serum INS, albumin, globulins and IGF-I, in addition to higher serum NEFA, urea and creatinine.

When energy intake is high, INS concentrations are also high, which promotes growth and/or BR recovery [1, 19, 22]. Such a positive correlation between energy intake and INS concentrations has been reported and this response was confirmed when lowering energy supply: concentrations of INS decreased during energy restriction [18].

### Adrenergic challenge

Our results regarding the individual responses to the ß-adrenergic challenge are in agreement with Chilliard et al. [13]. These authors found that basal NEFA and NEFA response to an isoprotenol (ISO) challenge with a similar dose to that used in our experiment (4 nmol/kg BW) were higher in underfed than overfed cows. Consistent with our findings, high correlations between the response area or maximal NEFA response and NEFA response at 15 min. (*r* = 0.95 and 0.98, respectively) were observed. A significant effect of BCS on the basal plasma level was also reported, thus concluding that NEFA response to ISO at 15 min. could provide an efficient method for *in vivo* studying the AT lipolytic potential. Our results, including trends of the response curves with regard to BCS groups, are very similar to those findings, except maximum value was obtained at 10 min. in our study. We also agree with the fact that maximal response occurred later when this response was higher which illustrates that lipolytic response to ISO take longer in underfed animals.

The significant correlation (*r* = 0.69) between plasma NEFA and NEFA maximal response to ISO confirms results obtained in lactating ewes by Bocquier et al. [23] and suggests that the adrenergic component of the lipolytic cascade plays a significant role in the regulation of basal plasma NEFA. The NEFA response to ISO challenge in that well-fed lactating ewes depended on body lipid mass but not on energy balance. By the contrary, in underfed ewes the NEFA response depended on energy balance and not on the body lipid mass. Adrenergic challenge was also useful in explaining the differences in interindividual adaptive strategies to underfeeding in the ewe. In underfed ewes Bocquier et al. [23] observed that the relative variation in milk yield was negatively correlated to NEFA+10 (*r* = -0.51), which show that ability to support lactation was related to the ability to mobilise body lipids.

### Potential contribution for a simplified method helping to identify individual adaptive capacities or robustness (intraflock variability)

There is evidence of the great potential of plasma NEFA as a powerful predictor of the nutritional status of the ruminant under determined circumstances. This parameter provides reliable information on the stage of the BR mobilization of the animal under exigent physiological status and/or when facing the consequences of being reared in fluctuating environments [3, 4]. The NEFA is thus recommended as a good diagnostic tool for health or reproductive interpretations, and we think that it could also be considered as a pertinent variable to be included in models aiming to analyze metabolic plasticity of ruminants when facing variability in feed availability and quality in a given timespan (i.e. individual robustness).

In previous works carried out by our team and aiming to characterize the energy metabolism of ewes in a typical round productive year, such a NEFA potential for illustrating the dynamic of individual BR status was confirmed in Romane [3] and Lacaune [4] meat and dairy ewes, respectively.

In the present study we evaluated the *in vivo* method with a β-adrenergic challenge. We confirmed our hypothesis that ewes with different, contrasted BCS would respond differently to a β-adrenergic challenge and that this response could be predicted at a given point (10 min.) of the plasma NEFA kinetic after the challenge, in function of the relationships between the different parameters responses at different times.

Chilliard et al. [13] using the same method, looked for a simplified procedure for predicting the lipolytic response curve with a smaller number of samples after ISO challenge. Consistent with our results (Figure 5, 6), they obtained a very good prediction of AUC from time 0 to 60 min either by the partial AUC from time 0 to 20 min (*r* = 0.95) or by the sole response at 15 (*r* = 0.95) or 20 min (*r* = 0.97). Measuring plasma NEFA in blood samples taken just before and at 15 (or 20) min after an ISO injection was thus considered as an efficient and simple way of predicting the maximal NEFA response of an individual and an AUC equivalent to one hour of sampling. Extra blood samplings at 5, 10, and 20 (or 15) min after ISO challenge only slightly increased the prediction of these parameters.

The adaptive capacity of a ruminant to an adrenergic challenge alias (i.e. pronounced energy shortage) is expressed by subsequent individual physiological responses at the short-, medium- and long-term timespans. Differences in the amplitude (gap between maximum NEFA response and basal NEFA), turnover (exponential slope when reducing plasma NEFA after maximal response) and length of those specific and combined processes are expected to be consistent with adaptive capacities’ differences between individuals reared under similar conditions.

A stronger lipolytic potential could be seen as a sight of the ultimate necessity of the animal to compensate their basic requirements by mainly using their BR. Indeed, when facing an undernutrition event (i.e. challenge), a higher and quicker BR mobilization peak (illustrated by plasma NEFA) could be a symptom of the incapacity of the animal to readjust its MER at the short-term. This would be in close relationship with their more or less efficient capacity of regulating (reducing) its feed intake and thus the individual MER, which would mean a higher dependency of their BR *per se* to cover energy requirements. Thus, under uniform conditions (i.e. same species, breed, physiological state, age, production system, feeding regimen…) less NEFA at a given point after the challenge would means that the animal is less depending from its BR. That individual, with more pronounced NEFA amplitude and quicker NEFA turnover would be, a priori, a better adapted animal when compared to its cohort. Such differences at the intragroup level were observed in our experiment (Figure 7). This could enable us to the potential effective use of this relative easy and quick method, for contributing to give useful information for identifying existing intraflock variability in individual robustness in practice at a given field situation.

## Conclusions

The findings confirmed the ability of these mature, dry, non-pregnant Mérinos d’Arles ewes to quickly overcome undernutrition situations by efficiently using their body reserves. The anabolic or catabolic responses to energy dietary manipulations were accompanied by synchronised metabolic regulation, resulting in differences in their metabolic and BCS profiles.

Because of the fact that lipolytic activity of adipose tissue differed among ewes with similar body condition status in the same group, our results also indicate the potential of using a simplified ß-adrenergic challenge protocol for identifying, at the intraflock level, individual differences in adaptive capacity to undernutrition.

## Acknowledgments

Authors are grateful to the technical staff of *Domaine du Merle* experimental farm, especially to Céline Maton, Jean-Dominique Guyonneau and Denis Montier for assisting with animal care and data collection.

## References

[1] Chilliard Y, Bocquier F, Doreau M. Digestive and metabolic adaptations of ruminants to undernutrition, and consequences on reproduction. Reprod Nutr Dev. 1998;38:131–52.

[2] Friggens NC, Brun-Lafleur L, Faverdin P, Sauvant D, Martin O. Advances in predicting nutrient partitioning in the dairy cow: recognizing the central role of genotype and its expression through time. Animal 2013; 7:89–101.

[3] González-García E, Gozzo de Figuereido V, Foulquie D, Jousserand E, Autran P, Camous S, et al. Circannual body reserve dynamics and metabolic profile changes in Romane ewes grazing on rangelands. Domest Anim Endocrinol 2014;46:37–48.

[4] González-García E, Tesniere A., Camous S, Bocquier F, Barillet F, Hassoun P. The effects of parity, litter size, physiological state, and milking frequency on the metabolic profile of Lacaune dairy ewes. Domest Anim Endocrinol 2015;50:32–44.

[5] Ferlay A, Chilliard Y. Effects of the infusion of non-selective β-, and selective β1-or β2-adrenergic agonists, on body fat mobilisation in underfed or overfed non-pregnant heifers. Reprod Nutr Dev. 1999;39:409–21.

[6] Teyssier J, Migaud M, Debus N, Maton C, Tillard E, Malpaux B, et al. Expression of seasonality in Merinos d’Arles ewes of different genotypes at the MT1 melatonin receptor gene. Animal 2011;5:329–36.

[7] Hanocq E, Bodin L, Thimonier J, Teyssier J, Malpaux B, Chemineau P. Genetic parameters of spontaneous spring ovulatory activity in Mérinos d’Arles sheep. Genet Sel Evol 1999;31:77–99.

[8] Debus N, Chavatte-Palmer P, Viudes G, Camous S, Roséfort A, Hassoun P. Maternal periconceptional undernutrition in Merinos d’Arles sheep: 1. Effects on pregnancy and reproduction results of dams and offspring growth performances. Theriogenology 2012;77:1453–65.

[9] INRA. Alimentation des bovins, ovins et caprins Besoins des animaux - Valeurs des aliments. Tables Inr. Paris, France: 2010.

[10] Russel AJF, Doney JM, Gunn RG. Subjective assessment of body fat in live sheep. J Agric Sci. 1969;72:451–4.

[11] Williamson DH, Mellanby JD. Hydroxybutyrate. In: Bergmeyer HB, editor. Methods of enzymatic analysis. New York Academic Press; 1974. p. 1836–9.

[12] Statistical Analysis Systems Institute (SAS). SAS language guide for personal computers. Cary, NC, USA: SAS Institute Inc; 2003. Version 9.3.

[13] Chilliard Y, Ferlay A, Desprès L, Bocquier F. Plasma non-esterified fatty acid response to a β-adrenergic challenge before or after feeding in energy underfed or overfed, dry or lactating cows. Anim Sci. 1998;67:213–23.

[14] Bocquier F, González-García E. Sustainability of ruminant agriculture in the new context: feeding strategies and features of animal adaptability into the necessary holistic approach. Animal 2011;4:7:1258–1273.

[15] Friggens NC, Ingvartsen KL, Emmans GC. Prediction of body lipid change in pregnancy and lactation. J Dairy Sci 2004;87:988–1000.

[16] Blanc F, Bocquier F, Agabriel J, D’hour P, Chilliard Y. Adaptive abilities of the females and sustainability of ruminant livestock systems. A review. Anim Res. 2006;55:489–510.

[17] Bauman DE, Currie WB. Partitioning of nutrients during pregnancy and lactation: a review of mechanisms involving homeostasis and homeorhesis. J Dairy Sci. 1980;63:1514–29.

[18] Cassady JM, Maddock TD, DiCostanzo A, Lamb GC. Initial body condition score affects hormone and metabolite response to nutritional restriction and repletion in yearling postpubertal beef heifers. J Anim Sci. 2009;87:2262–73.

[19] Sosa C, Abecia JA, Carriquiry M, Forcada F, Martin GB, Palacín I, et al. Early pregnancy alters the metabolic responses to restricted nutrition in sheep. Domest Anim Endocrinol. 2009;36:13–23.

[20] Caldeira RM, Portugal A V. Interrelationships between body condition and metabolic status in ewes. Small Rumin Res. 1991;6:15–24.

[21] Atti N, Bocquier F KG. Performance of the fat-tailed Barbarine sheep in its environment: adaptive capacity to alternation of underfeeding and re-feeding periods. A review. Anim Res. 2004;53:165–76.

[22] Delavaud C, Bocquier F, Baumont R, Chaillou E, Ban-Tokuda T, Chilliard Y. Body fat content and feeding level interact strongly in the short-and medium-term regulation of plasma leptin during underfeeding and re-feeding in adult sheep. Br J Nutr. 2007;98:106–15.

[23] Bocquier F, Ferlay A Chilliard Y. Effects of body lipids and energy balance on the response of plasma non-esterified fatty acids to a ß-adrenergic challenge in the lactating dairy ewe. In: McCracken KK, Unsworth EF, Wylie ARG, editors. Energy metabolism of farm animals. Proceedings of the 14th symposium on energy metabolism inform animals, Newcastle, Northern Ireland; 1998. 167–170. CAB International, University Press, Cambridge.

